## Supplementary Materials for "Single-nucleus RNA-sequencing in pre-cellularization *Drosophila melanogaster* embryos"

Ashley Albright[1], Michael Stadler[1], Michael Eisen[1,2]
[1] Department of Molecular and Cell Biology, University of California Berkeley, United States
[2] Howard Hughes Medical Institute, United States

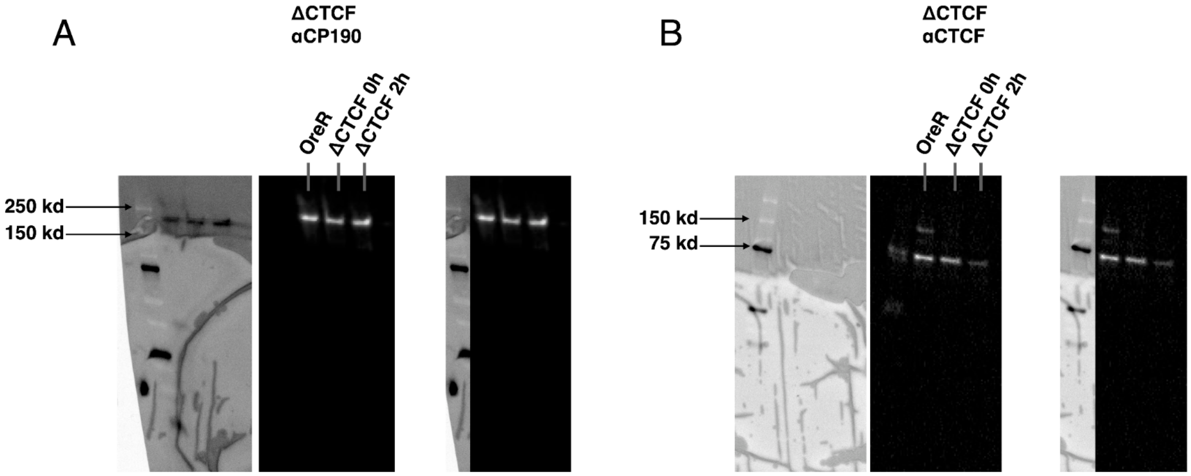


**Supplementary Figure 1:** (a) Western blotting of OreR, 0h and 2h *dCTCF*$^{mat-/-}$ embryos using an antibody to *Cp190,* another insulator protein, as a control. (b) Western blotting of OreR, 0h and 2h *dCTCF*$^{mat-/-}$ embryos using an antibody to *dCTCF.* A cross-reactive band appears at approximately 75 kd, with the *dCTCF* band appearing at approximately 130 kd.

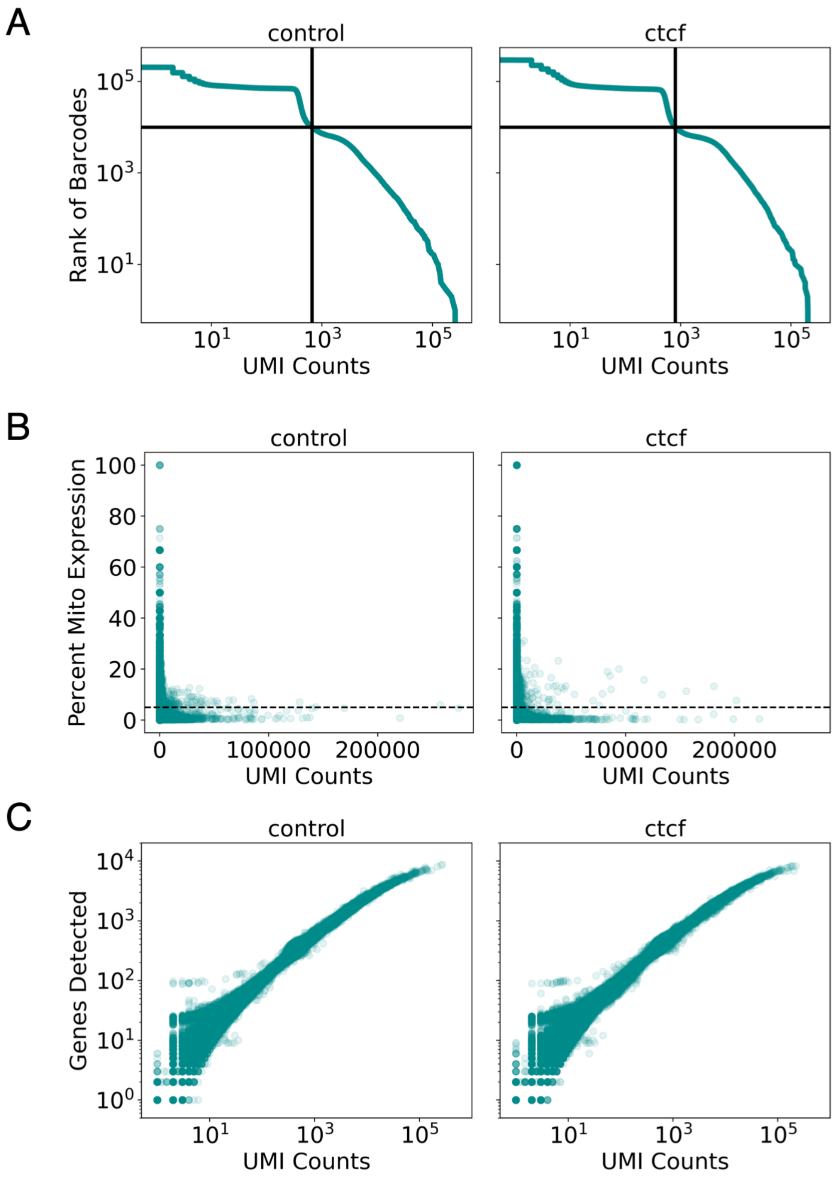


**Supplementary Figure 2:** (a) Knee plot for barcodes ranked by the number of UMIs versus UMI counts for control (left) and *dCTCF*$^{mat-/-}$ (right) experiments. Black line indicates the position of the 10,000$^{th}$ (expected number of cells) on each axis. (b) Percent mitochondrial expression per nucleus in control (left) and *dCTCF*$^{mat-/-}$(right) nuclei. Dashed line represents 5% mitochondrial expression, or the cutoff used for filtering the data. (c) Number of genes detected per nucleus by UMI counts in control (left) and *dCTCF*$^{mat-/-}$(right) nuclei.

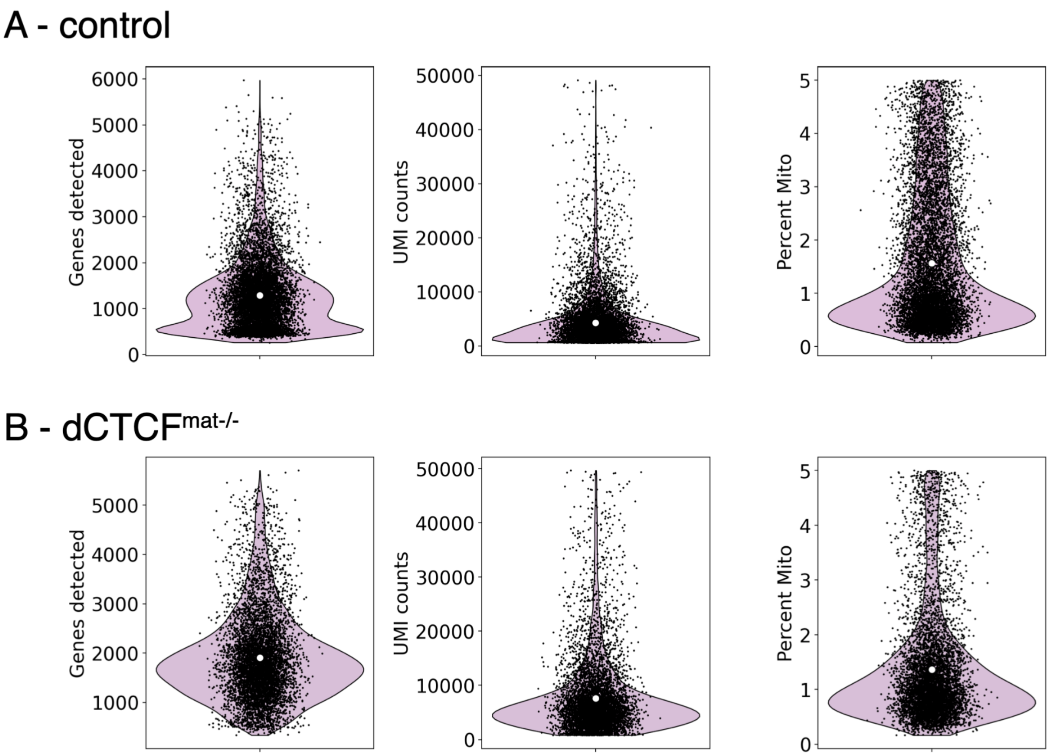


**Supplementary Figure 3:** Number of genes detected (left), UMI counts (middle), percent mitochondrial expression (right) per nucleus after filtering in (a) control and (b) *dCTCF$^{mat-/-}$* experiments.

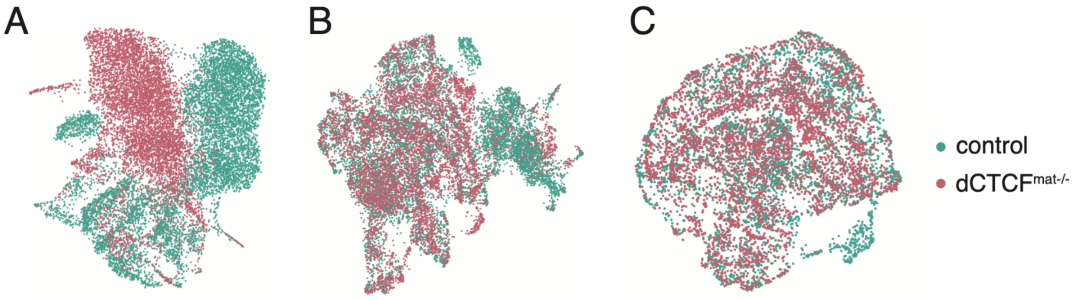


**Supplementary Figure 4:** Two-dimensional UMAP embedding of control (teal) and *dCTCF$^{mat-/-}$* (pink) nuclei (a) before and (b) after batch correction using scVI.

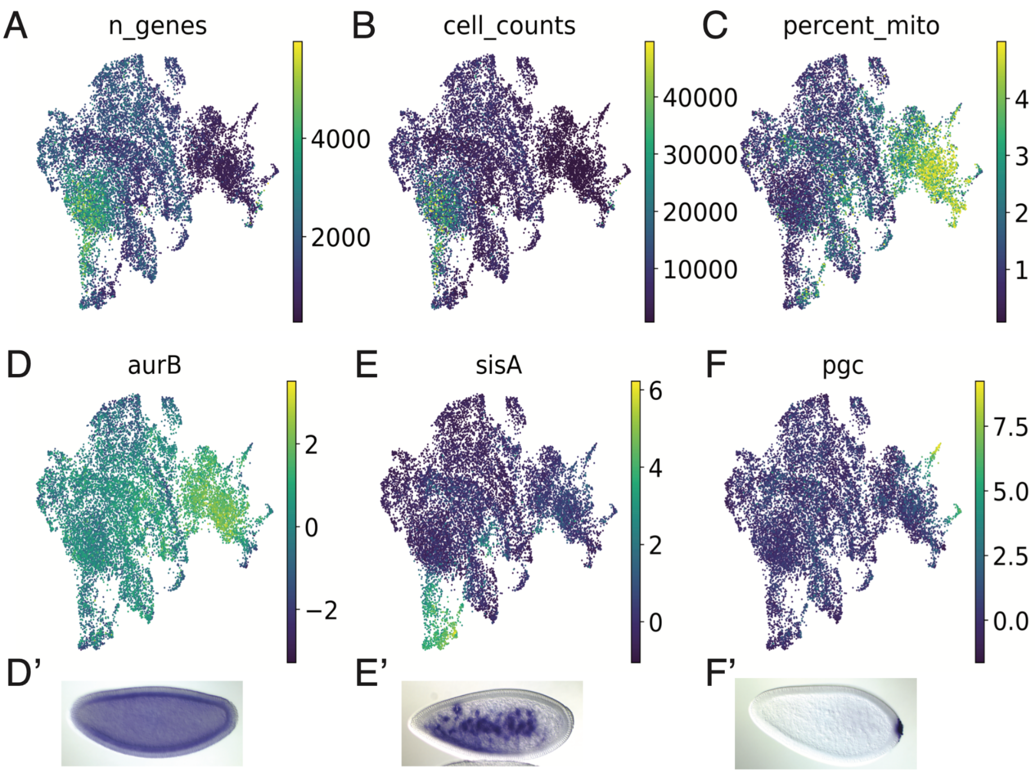


**Supplementary Figure 5:** Two-dimensional UMAP embedding of nuclei before additional filtering colored by (a) number of genes detected, (b) UMI counts, (c) percent mitochondrial expression. (d-f) log(scvi normalized expression) of three genes with representative *in situ* hybridizations below for (d) cell cycle gene *aurB*, (e) yolk nucleus marker, *sisA* (f) and pole cell marker *pgc*.

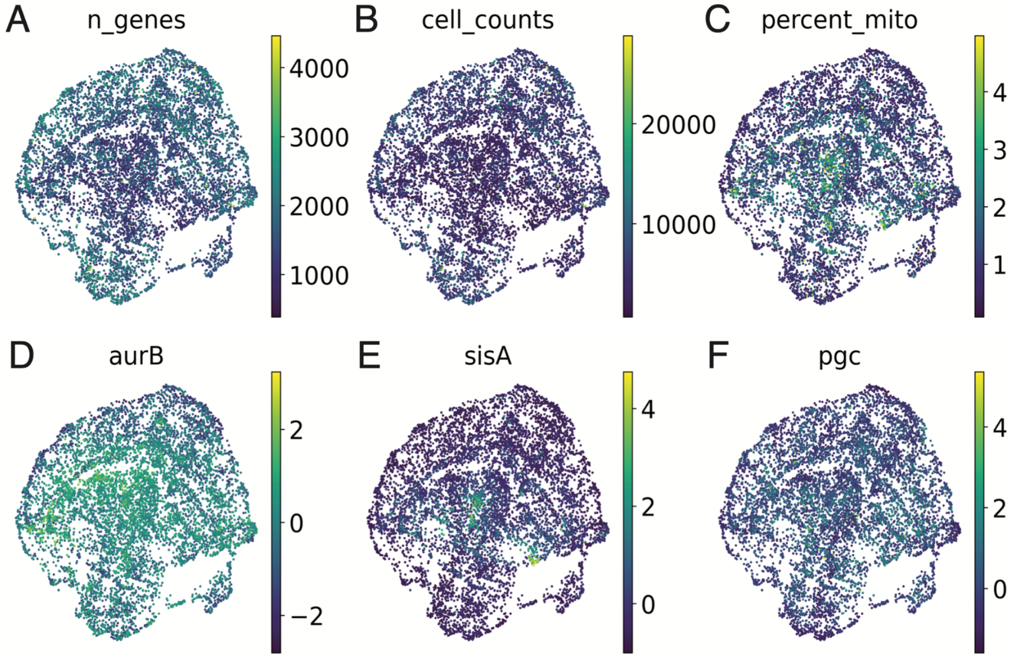


**Supplementary Figure 6:** Two-dimensional UMAP embedding of nuclei after removal of clusters with high percent mitochondrial expression, *aurB* expression, *sisA* expression, and *pgc* expression colored by (a) number of genes detected, (b) UMI counts, (c) percent mitochondrial expression. (d-f) log(scvi normalized expression) of three genes with representative *in situ* hybridizations below for (d) cell cycle gene *aurB*, (e) yolk nucleus marker, *sisA* (f) and pole cell marker *pgc*.

| 0 | 1 | 2 | 3 | 4 | 5 | 6 | 7 | 8 | 9 |
|---|---|---|---|---|---|---|---|---|---|
| CREG | ths | CG3502 | GstD10 | htl | fkh | G6P | cv-2 | lncRNA: CR43685 | CG2233 |
| CG14561 | pyr | CG14204 | CG32052 | CadN | lncRNA: CR43126 | lncRNA: CR46455 | Sarm | CG3408 | Cht7 |
| CG34214 | Trim9 | Lgr1 | CG15128 | if | tll | hth | net | CG2225 | CG32365 |
| Mco1 | wb | CG42342 | Ae2 | CG9005 | out | Meltrin | Samuel | CG43232 | CG7896 |
| CG5888 | CG46458 | lncRNA: CR44929 | Ilp4 | CG33725 | disco | Pgd | CG31523 | BI-1 | Pld |
| Cpr31A | mfas | frma | spg | kibra | CG44837 | path | mtd | CG31300 | Gad1 |
| mnd | Tet | lncRNA: CR44732 | CG9775 | Mid1 | byn | jbug | CG45263 | fusl | Proc-R |
| CG4702 | raskol | CG14205 | CG6398 | stumps | srp | lncRNA: CR46454 | CheA84a | mtDNA-helicase | Ca-Ma2d |
| geko | CG12038 | Damm | asRNA: CR44095 | CG10663 | bbg | CG3764 | heph | CG14931 | CG2652 |
| asRNA: CR44095 | Antp | CG7900 | lncRNA: CR44697 | lncRNA: CR45361 | CG15544 | Wdr62 | mew | CG44836 | EndoGI |
| CG13868 | Meltrin | nrm | CG4570 | msn | rib | CG42851 | CG13654 | ClC-b | CCKLR-17 D3 |
| upd1 | S-Lap8 | Adgf-A | CG13033 | NetA | NK7.1 | blot | CG8834 | CG14864 | CG15673 |
| CG13827 | IFT46 | lncRNA: CR44526 | hrg | pigs | CG2930 | D2hgdh | ush | GstD8 | hebe |
| halo | CG17834 | otk | Hf | Jhedup | hkb | raskol | Cyp313b1 | lncRNA: CR44365 | CG15239 |
| CG15382 | dally | Bili | Ady43A | Nplp1 | Fili | aop | CG12934 | Hsp67Bc | CG31675 |
| D | Wdr62 | CG7886 | NetA | T48 | Obp56d | prd | Cip4 | CG42266 | CG17321 |
| MsrA | mtgo | Dhc64C | neur | CG11357 | CG1090 | CG5043 | NK7.1 | ND-B17.2 | org-1 |
| esn | boi | noc | Elba3 | for | mthl3 | salm | CG44085 | CG31817 | Gs2 |
| Acer | CG9257 | CG15611 | IM14 | CG12177 | alpha-Est1 | CG15628 | CG2016 | Drp1 | CG6959 |
| PQBP1 | CG14618 | CG10553 | edl | CG5004 | nab | jigr1 | Ubx | CG5027 | Fcp3C |

**Supplementary Table 1:** Top 20 marker genes representing each cluster as determined by sc.tl.rank_genes_groups.

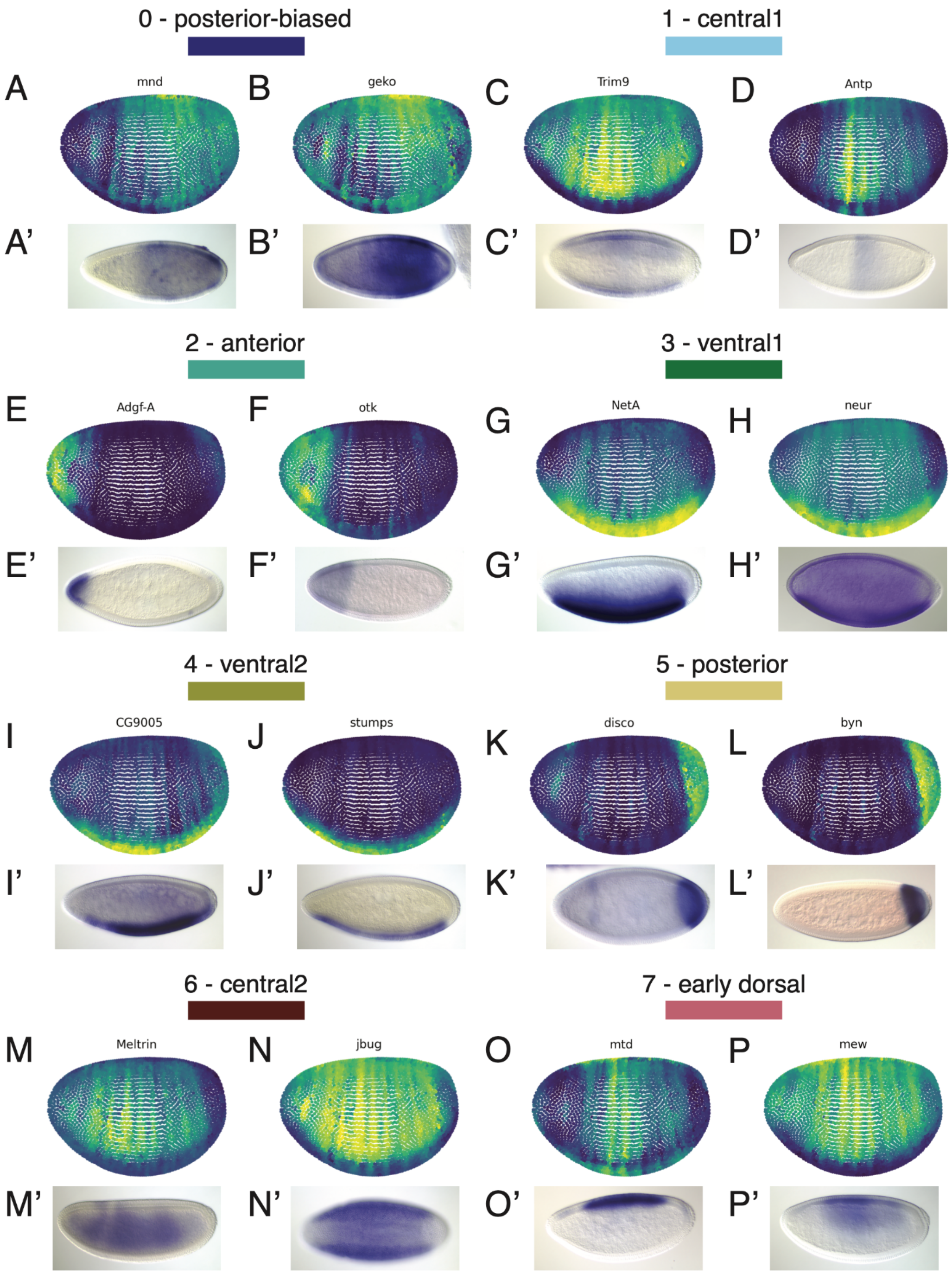


**Supplementary Figure 7:** (a-p) Representative virtual (top, a-p) and Berkeley Drosophila Genome Project (bottom, a'-p') *in situ* hybridizations for additional marker gene expression within each cluster as indicated. This supplemental figure accompanies Figure 2 in the main text.

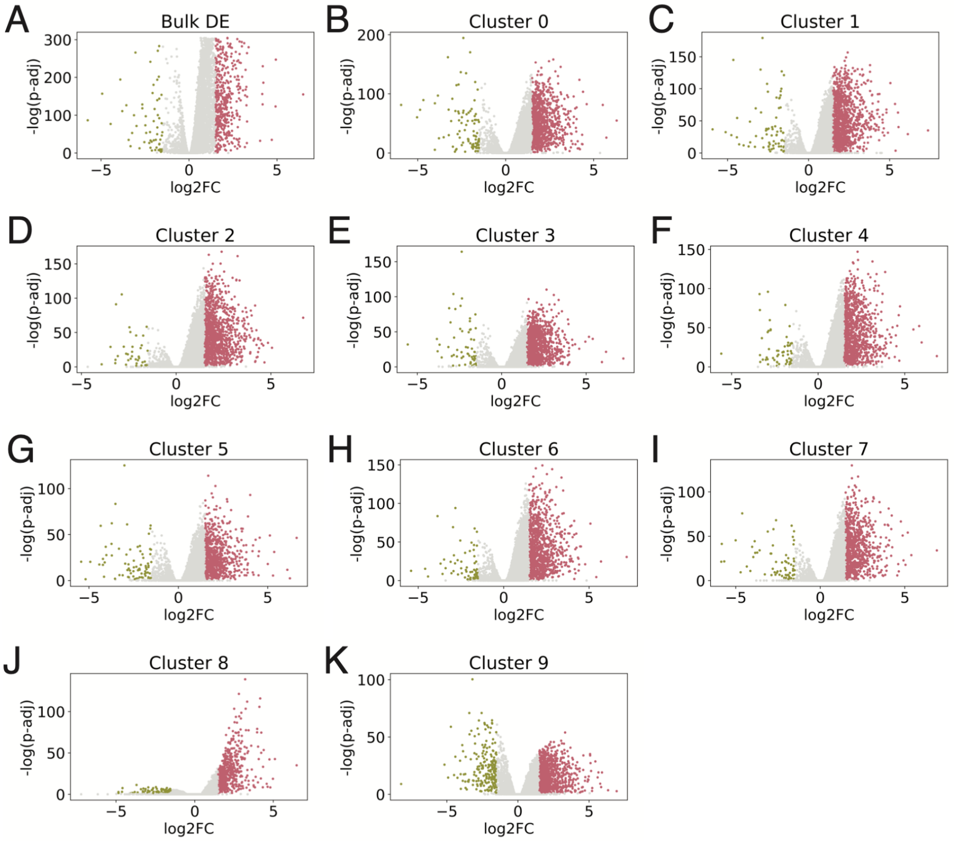


**Supplementary Figure 8:** Volcano plots of log2FC (log2(fold-change)) by the log of the adjusted p-value (p-adj) for differential expression calculated in bulk (top middle) and in individual clusters as indicated. Colored dots indicate genes with significant differential expression, an absolute value of log2FC >= 1.5 and p-adj < 0.05. Significantly down-regulated genes are indicated in green, significantly up-regulated genes in pink, and non-significantly differentially expressed genes in light gray.

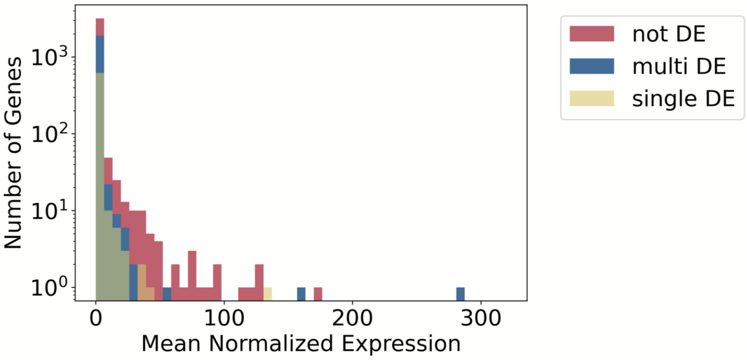

**Supplementary Figure 9:** Histogram of average gene expression of differentially expressed genes in one cluster (yellow), differentially expressed in multiple clusters and/or in bulk (blue), and non-differentially expressed genes (red). Each count on the y-axis represents a single gene.
